# A general substitution matrix for structural phylogenetics

**DOI:** 10.1101/2024.09.19.613819

**Authors:** Sriram G Garg, Georg KA Hochberg

## Abstract

Sequence-based maximum likelihood (ML) phylogenetics is a widely used method for inferring evolutionary relationships, which has illuminated the evolutionary histories of proteins and the organisms that harbour them. But modern implementations with sophisticated models of sequence evolution struggle to resolve deep evolutionary relationships, which can be obscured by excessive sequence divergence and substitution saturation. Structural phylogenetics has emerged as a promising alternative, because protein structure evolves much more slowly than protein sequences. Recent developments protein structure prediction using AI have made it possible to predict protein structures for entire protein families, and then to translate these structures into a sequence representation - the 3Di structural alphabet - that can in theory be directly fed into existing sequence based phylogenetic software. To unlock the full potential of this idea, however, requires the inference of a general substitution matrix for structural phylogenetics, which has so far been missing. Here we infer this matrix from large datasets of protein structures and show that it results in a better fit to empirical datasets that previous approaches. We then use this matrix to re-visit the question of the root of the tree of life. Using structural phylogenies of universal paralogs, we provide the first unambiguous evidence for a root between and archaea and bacteria. Finally, we discuss some practical and conceptual limitations of structural phylogenetics. Our 3Di substitution matrix provides a starting point for revisiting many deep phylogenetic problems that have so far been extremely difficult to solve.

## Introduction

The field of phylogenetics has evolved from relying on morphological comparisons to sophisticated sequence-based analyses (Whelan et al., 2001). The advent of computational methods marked a turning point, introducing a range of algorithms from Neighbour-Joining (NJ) (Saitou and Nei, 1987) and Maximum Parsimony (MP) (Farris, 1970; Fitch 1971) to Maximum Likelihood (ML) (Felsenstein, 1981) and Bayesian inferences (Rannala and Yang, 1996; Mau and Newton, 1997) on nucleotide and amino acid sequences. Each methodological leap has brought with it a deeper understanding of evolutionary history through better trees. Among the various phylogenetic methods, Maximum Likelihood (ML) approaches have emerged as particularly powerful tools for modelling evolutionary processes (Posada and Crandall, 2021). The flexibility and robustness of ML techniques have made them indispensable for contemporary phylogenetic studies, especially those tackling large datasets or seeking to resolve deep evolutionary relationships. But especially deep, sequenced-based phylogenetics remains difficult. Substitution saturation is a particular challenge, in which each site in the alignment has accumulated multiple substitutions over a branch of interest (Brown, 1982, Phillippe and Forterre 1999). Depending on the accuracy of the substitution model of sequence evolution, saturation can lead to spurious phylogenetic signals and artefacts in phylogenetic trees (Felsenstein, 2003). The problem of saturation cannot always be solved by adding more sequences (Philippe et al., 2011) or better models of sequence evolution.

Saturation is a relevant problem for the identification of the root of the tree of life. It is traditionally placed on the branch between bacteria and archaea (Gouy et al., 2015), which has important implications for the nature of the Last Universal Common Ancestor (LUCA). This inference is based on paralog rooting with universally duplicated genes, where the paralogs reciprocally root each other (Iwabe et al., 1989). Although this root is tacitly accepted by the majority of biologists, the paralog trees it is based on are riddled with potential problems. In all previous attempts, the branch between universal paralogs remains so long as to be probably saturated (Brown and Doolittle, 1995; Philippe and Forterre, 1999; Gouy et al., 2015; Mahendrarajah et al., 2023). This means that the root position within each paralog might be mostly determined by the preferences of the substitution model, rather than real phylogenetic signal, which has been erased almost entirely. Some phylogenetics therefore still consider the root of the tree of life an unsolved problem (Gouy et al., 2015).

Structural phylogenetics offers a potentially powerful alternative to traditional sequence-based approaches. Structures evolve much more slowly than sequences, and if a model for structural evolution could be inferred, this could help resolve phylogenies that are beyond the reach of sequenced-based methods. Early attempts at this idea were limited by the lack of high-quality protein structures or reliable methods of scoring multiple sequence alignments of protein structures (Johnson, Šali, et al., 1990; Johnson et al., 1990; Balaji et al., 2001; Balaji and Srinivasan, 2001). This changed with the advent of artificial intelligence models than can predict protein structures with good accuracy (Jumper et al., 2021; Varadi et al., 2023). The availability of a large database of structures has prompted researchers to mould this novel source of information for identification of structural homologs in a process similar to BLAST. Chief among these tools is FoldSeek which translates the 3D information in predicted and experimentally determined structures into 20 unique characters the authors call the 3Di alphabet (Kempen et al., 2023). The advantage of using an alphabet of 20 characters is that it enables the direct use of these 3Di characters in conventional implementations of amino acid-based likelihood methods.

The conversion of a large dataset of 3D structures into the 3Di alphabet allows the computation of a scoring matrix like the BLOSUM scoring matrix commonly employed by Multiple Sequence Alignment programs (Kempen et al., 2023). This scoring matrix enables the quick identification of structural homologs of proteins which has been very successful in the identification of divergent orthologs. Such a scoring system also allows to compute a similarity score (*fident* in case of FoldTree) which can then be used to compute Neighbour Joining (NJ) trees as demonstrated by FoldTree (Moi et al., 2023). Furthermore, one could also calculate a substitution matrix from this BLOSUM style scoring matrix which can be directly implemented in ML approaches such as in the case of 3DiPhy (Puente-Lelievre et al., 2024). Neither of these approaches correspond to standard maximum likelihood phylogenetics for amino acids: FoldTree’s neighbour joining method is fast and simple but inherits all limitations of classical neighbour joining in that it relies on the true distance between sequences being close to their observed distance (an assumption that is often violated in realistic datasets) and it does not account for among site rate variation (Mihaescu et al., 2009). 3DiPhy does use a full likelihood model, which can account for these phenomena however, its substitution matrix is derived from a BLOSUM-like alignment scoring matrix. Such matrices are constructed by counting co-occurrences of particular characters in sequence pairs, rather than inferring their contents using maximum likelihood (Le and Gascuel, 2008). In standard sequence phylogenetics the BLOSUM matrix has long been superseded by empirical models which are inferred in a full phylogenetic likelihood framework, and generally result in a much better fit to empirical data (Le and Gascuel, 2008).

These features of existing structural phylogenetics frameworks motivated us to infer a new substitution model using a phylogenetic maximum likelihood framework. This substitution model can in theory be directly inferred from each alignment in the form of a General Time Reversible (GTR) model but inferring a substitution matrix for a 20-letter alphabet from a single multiple sequence alignment is difficult and prone to overfitting. For conventional protein models, this problem is solved by combining large numbers of protein alignments and inferring from them one substitution model that best describes all the data. Once computed, this general model, also denoted as *Q,* can then be used for individual protein families, which avoids overfitting using GTR. Here we make use of AlphaFold and a recently developed protein large language model to infer a general substitution matrix for structural phylogenetics. We show that this Q-matrix outperforms all previous methods to use 3Di characters to infer ML phylogenies. Finally, we use our Q-matrix to re-infer the phylogenies of universal paralogs and photosystems to settle long-standing questions in deep evolution that previously suffered from saturation.

## Results and Discussion

### Estimation of the 3Di Q-matrix

We set out to compute a general Q-matrix for structural phylogenetics. Given a large enough dataset, this is straightforward to achieve using the QMaker routine of IQ tree (Minh et al., 2021). We used two strategies to gather a large dataset of protein families and their predicted structures. Our first goal was to use the set of 6653 protein families that was used to infer a Q matrix in the initial study by Minh *et al*. To avoid having to predict AlphaFold models for every sequence in this large database, we opted to use a recently developed bilingual large language model, ProtT5. This model was trained to directly translate between an amino acid sequence and its corresponding 3Di sequence, without having to infer an AlphaFold model (Heinzinger et al., 2023). We used this method to translate all sequences in the PFAM dataset from AA-sequences to 3Di. The ProtT5 model is not perfect, as it introduces some randomness into the 3Di translation, meaning that translating to a 3Di sequence from the same input amino acid sequence results in a slightly different prediction (Supplementary Figure 1A-C). In addition, when comparing 3Di sequences extracted from AlphFold2 structures to the same 3Di sequence predicted with the LLM model, we found large numbers of sequences in which the AlphaFold and LLM predictions had low pairwise identities (Supplementary Figure 1D-F, Supplementary Figure 2). In order to safeguard against potential errors in estimating the substitution model using incorrectly translated 3Di sequences we also estimated a separate Q matrix using 3Di sequences extracted from AlphaFold predictions. We employed FoldSeek to cluster the SwissProt AlphaFold Database. These 1660 AF clusters (hereafter AF-db) were used along with 3Di translation of the 6653 protein families (hereafter Pfam-db), for the QMaker pipeline. Crucially, both sets of 3Di sequences were then aligned using the alignment program *mafft* using the 3Di scoring matrix from (Kempen et al., 2023) instead of the standard BLOSUM62 matrix used for amino acid alignments.

We then estimated a tree for each of 3Di Multiple Sequence Alignments (MSAs) in our two datasets using the GTR20 model despite the concern of model overfitting given the unique nature of the 3Di alphabet and the lack of other models that could serve as the initial starting model. These initial trees were then used to estimate a single Q-matrix that best explains the respective sets of MSAs as described in the QMaker pipeline (Minh et al., 2021). This resulted in two Q-matrixes hereafter denoted as Q.3Di.AF and Q.3Di.LLM. The two Q-matrices estimated were very similar with minor differences in exchangeabilities (Figure 1C) with a Pearsons correlation of 1 (Figure 1D). We then checked if these matrices are preferred by IQ-Tree’s *modelfinder* over the GTR20 or the previously published 3DiPhy model using a test set of 6653 3Di MSAs from PFAM that were not used for estimating the Q-matrix. Indeed, the 3DiPhy model is only preferred in 278 MSAs over 6267 MSAs that prefer either the Q.3Di.AF or the Q.3Di.LLM model, which are practically the same (Figure 1B-D). This increased our confidence that we had successfully captured the mechanism of change describing the mutability in the structural alphabet across a wide range of proteins. In the analyses of specific protein families that follow, IQ-Tree’s *modelfinder* predominantly chose Q.3Di.AF over Q.3Di.LLM or GTR20 according to the Corrected Akaike Criterion (AICc). Generally, we encourage future users of these matrices to always test if using Q.3Di.AF changes any conclusions in cases where Q.3Di.LLM is the better fit model. This is because the AF matrix is much less affected by the misprediction issues than the LLM (which we discuss further below).

**Figure 1:**
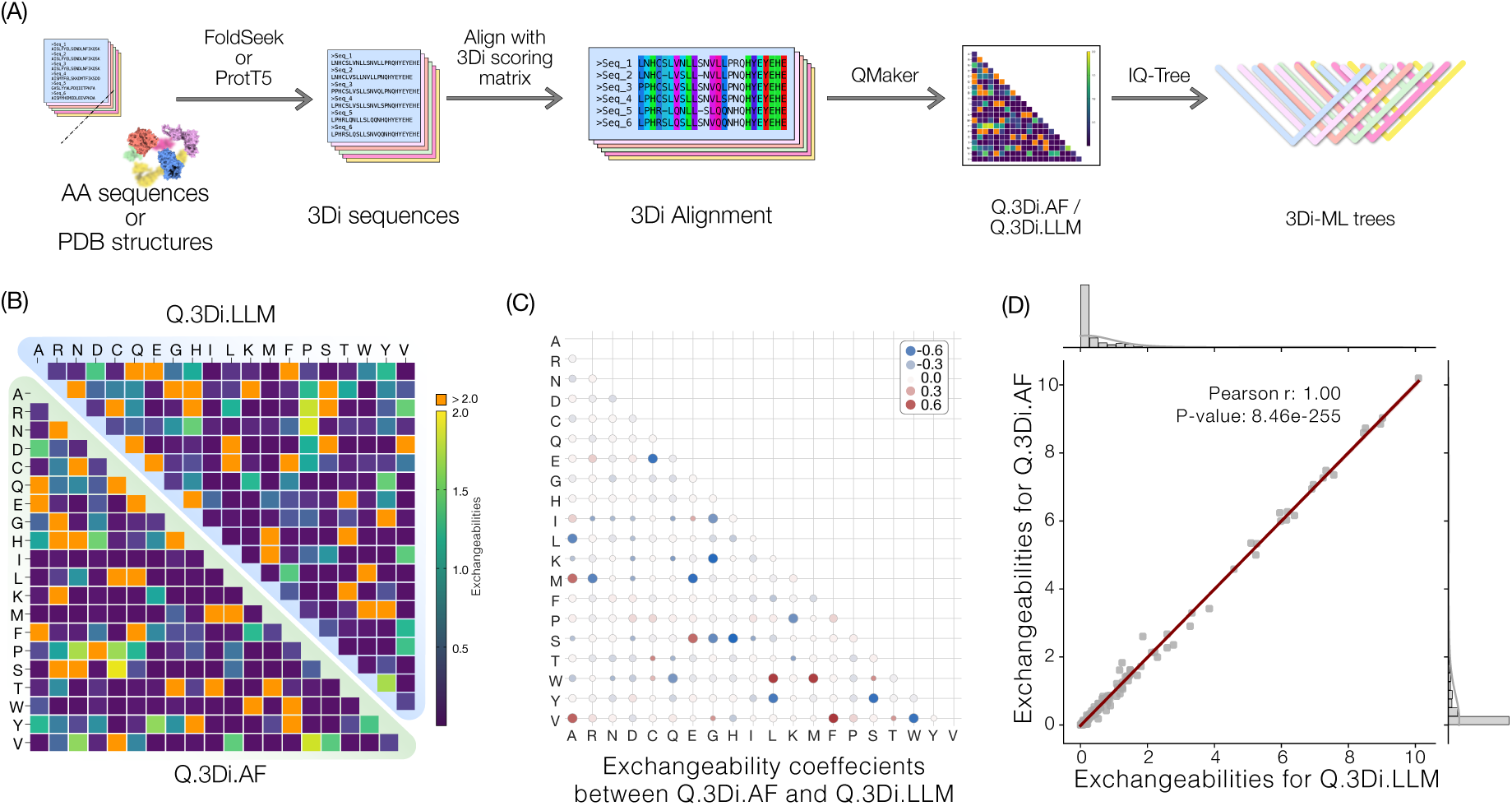
(A) Overview of the pipeline employed in the manuscript. Briefly, Amino acid (AA) or PDB structures were translated into 3Di characters using FoldSeek or the bilingual ProtT5 model. These 3Di characters are aligned with MAFFT using the 3Di scoring matrix before being used to estimate the general substitution Q-matrix using QMaker which were subsequently used to estimate 3Di-ML trees using IQ-tree. (B) Lower triangular portion is a representation of the Q-matrix estimated from 1660 AF clusters while the upper triangular section denotes the Q matrix estimated from 6653 PFAM clusters translated to 3Di alphabet using the ProtT5 bilingual language model. In both cases values higher than 2 are coloured orange. (C) Ratio of exchangeabilities between the Q.3Di.AF and the Q.3Di.LLM matrix. Each square represents the value 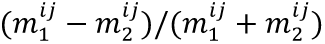 where m1 and m2 represent Q.3Di.AF and Q.3Di.LLM respectively. (D) Pearsons correlation between the exchangeabilities of the two matrices indicating very little differences between the two matrices

### Rooting the ToL using structural phylogenetics

Rooting the tree of life is a particularly challenging problem owing to the lack of outgroups that can reliably root phylogenetic trees. Paralog rooting is a powerful method which uses phylogenetic trees with duplicated genes that reciprocally root each other. In most cases the paralogs root each other along the same branch recovering an unambiguous root for the species tree containing the paralogs. However, in cases of highly divergent paralogs, the two paralogs sometimes do not agree on the same root (Figure 2A). We tested if our new matrix can help improve trees used to root the tree of life using two universal paralogs that have been previously used for this purpose: Elongation factors and catalytic and non-catalytic subunits of the rotary ATPase. We begin with the Elongation factor phylogeny. Elongation factor EF-Tu/EF-2 delivers aminoacyl-tRNAs to the A-site of the ribosome while the Elongation Factor EF-G/EF-1A catalyses the translocation of the peptidyl-tRNA (Miller, 1972). Both paralogs are conserved across the tree of life, making them an ideal candidate for paralog rooting (Baldauf et al., 1996; Philippe and Forterre, 1999; Gouy et al., 2015). In all previous attempts to root the tree of life using EF-G and EF-Tu, the branch separating the paralogs is extremely long and potentially completely saturated, which implies that the position of the root within each paralog might be determined entirely by the substitution model and not by any synapomorphies between the paralogs. In addition, the two paralogs do not root each other consistently increasing the uncertainty.

**Figure 2:**
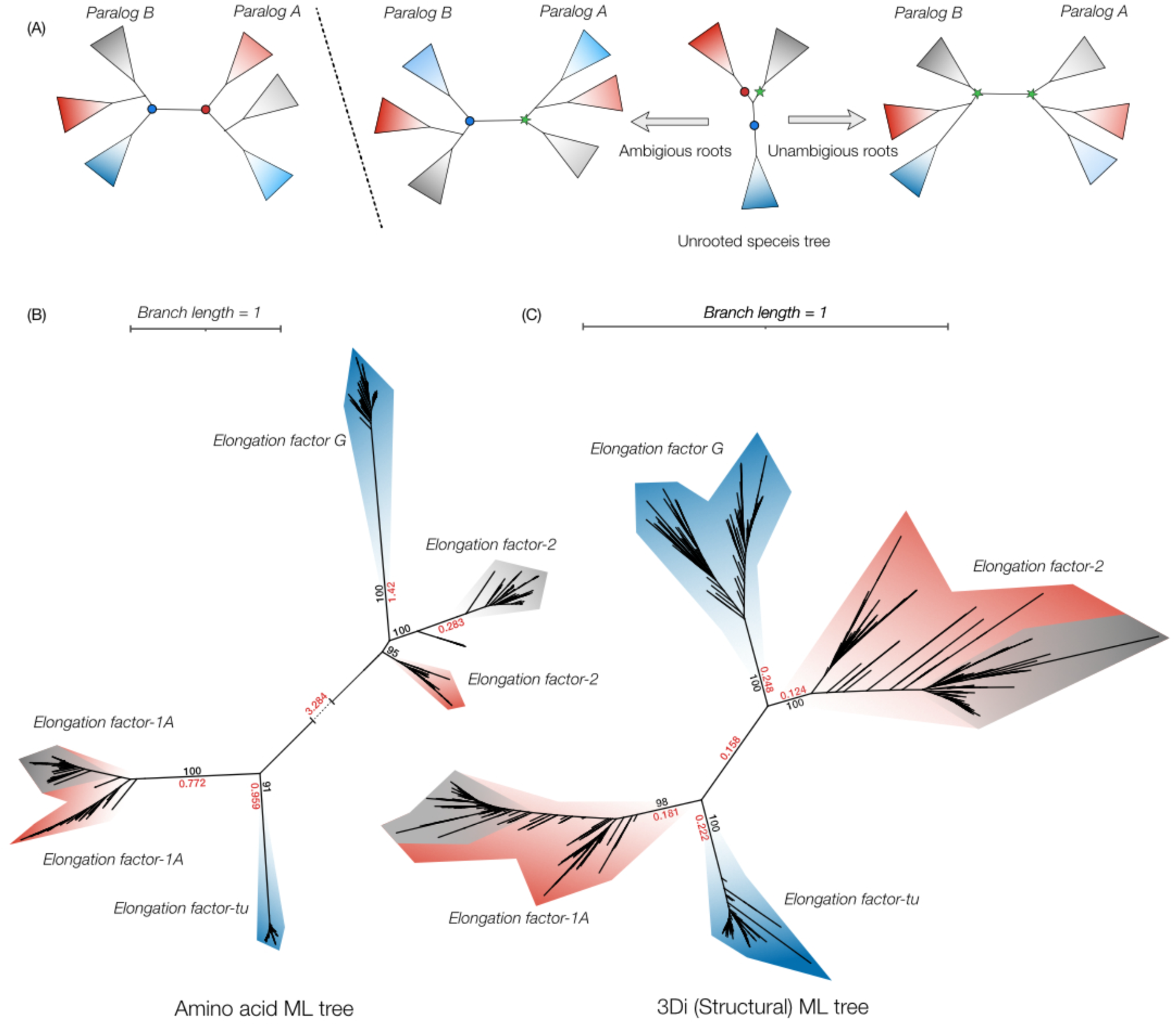
(A) A schematic representation of paralog rooting. Three possible root positions are shown with the “true” root depicted with a green star and two other possible roots with circles. In the scenario where paralogous rooting is successful both paralog subtrees reciprocally root each other (right). Other possible scenarios are also shown where the paralog subtrees are ambiguously rooted (left). (B) Amino acid ML tree containing 1076 EF-tu and EF-G homologs from eukaryotes, bacteria and archaea. The mitochondrial and plastid encoded copies are not included. Note that the branch separating the EF-tu and EF-G is broken for illustration. (C) 3Di structural ML tree estimated 3Di sequences and the Q.3Di.AF model from the predicted AlphaFold structures of 1069 EF-tu and EF-G homologs. In both cases blue, red and grey clades represent bacteria, archaea and eukaryotes respectively. Numbers in red, black indicate branch lengths and ultrafast bootstrap supports respectively.

To test if our new matrix can help solve this problem, we first assembled a dataset of 1076 homologs of EF-Tu and EF-G. In an amino acid-based ML tree we also recover a very long branch (Branch length (BL) = 3.284) between the two paralogs albeit still separating the bacteria and archaea (Figure 2A). In line with previous phylogenies, this tree recovers different roots for the tree of life in the two paralogs: between bacteria and archaea plus eukaryotes, and between archaea and bacteria plus eukaryotes (Figure 2B). We then extracted 3Di sequences from 1076 AlphaFold predictions using FoldSeek (see methods) and utilized our new Q.3Di.AF Q-matrix as the substitution model, to estimate a new tree of the EF-G and EF-tu paralogs. This recovered a phylogenetic tree with the length of the branch separating the paralogs far below 1 (0.186). Crucially, the root position is now consistent in both the paralogs and indicates a root between archaea and bacteria for life (Figure 2C). The archaea in both paralogs remain paraphyletic, which is consistent with the two-domain tree of life.

Another universally conserved paralogous gene family used to root the tree of life are the catalytic and non-catalytic subunits of the rotary ATPase. The head group of the rotary ATPase is a hexamer consisting of two subunits, only one of which is catalytic (Figure 3A). The bacterial and mitochondrial ATPases are called the F_0_F_1_-ATPases, and their subunits are called F1-alpha and F1-beta for the non-catalytic and catalytic subunits respectively (Grüber et al., 2001). The archaeal ATPase is called the V-ATPase and shares a similar architecture with a non-catalytic and a catalytic subunit in its headgroup (Figure3A). Owing to the endosymbiotic event between archaea and bacteria at eukaryogenesis, the eukaryotes and archaea also share this ATPase which in eukaryotes is in the vacuole, where it functions to acidify lysosomes (Gogarten et al., 1989). The archaeal/eukaryotic subunits are named V1-beta and V1-alpha for the non-catalytic and catalytic subunits respectively (Grüber et al., 2001; Cross and Müller 2004). A recent analysis on rooting the ToL using the ATPase subunits (Mahendrarajah et al., 2023) recovers a tree that separates the four major subunits with extremely long basal branches (Figure 3B). This tree is consistent with the idea that the catalytic and non-catalytic subunits originated before the divergence of archaea and bacteria, and roots the tree of life between these two domains. The same study also identified an early transfer of the archaeal non-catalytic subunit into bacteria, however, the catalytic counterpart to this transfer was not recovered in the catalytic sub-tree suggesting multiple transfer events (Figure 3B).

**Figure 3:**
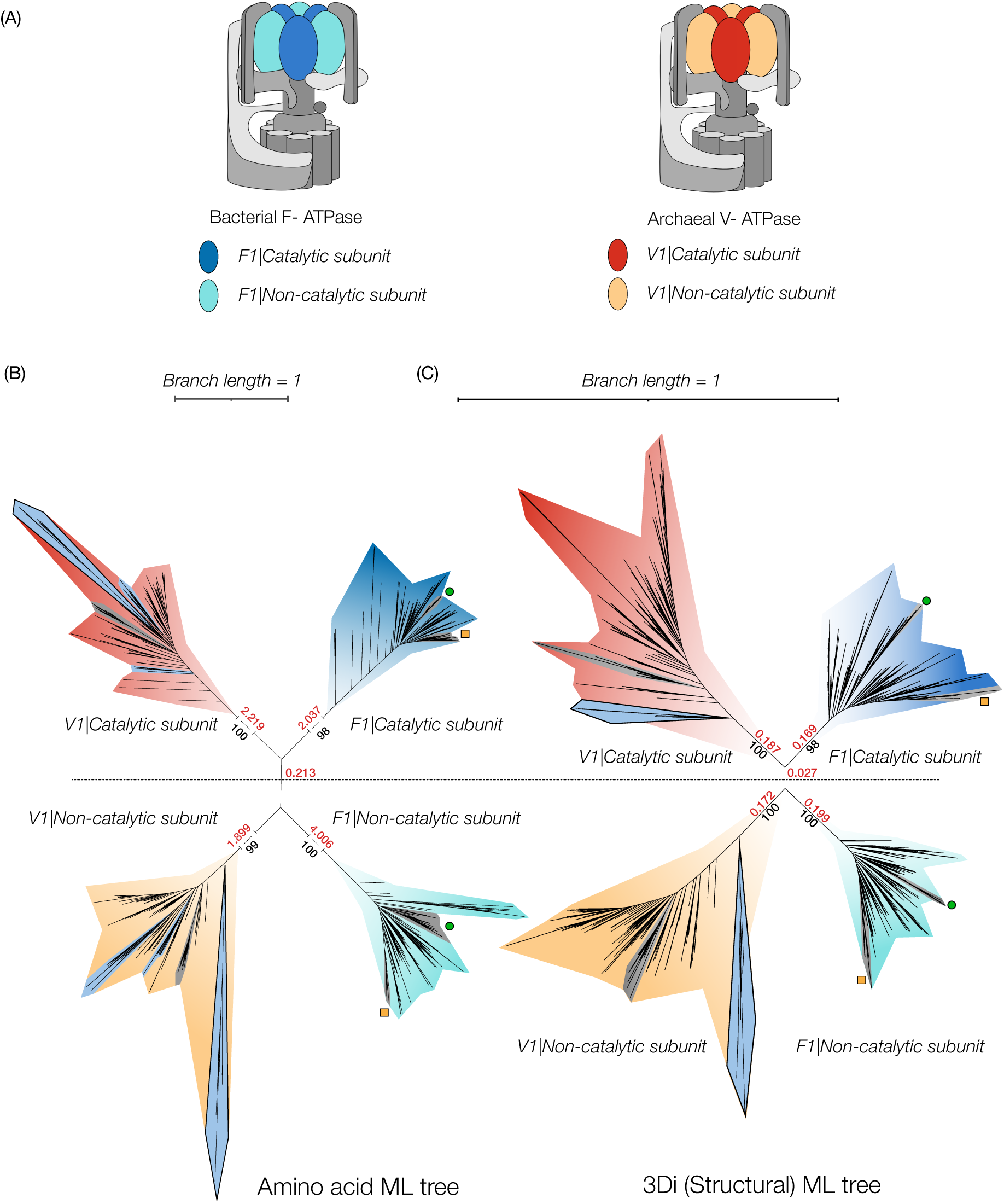
(A) A schematic representation of the bacterial and archaeal ATPase highlighting the subunits under investigation. They are represented using the same colours in the phylogenetic trees. (B) Amino acid ML tree of 1520 sequences across the ToL reproduced from Mahendrarajah et al., 2023 of the catalytic and non-catalytic subunits of bacterial, archaeal, and eukaryotic rotary ATPase. The early branching transfer from bacteria and archaea in the non-catalytic V1 clade is highlighted in blue with a black outline. The corresponding clade in the V1 catalytic clade branches deep inside of the archaeal sequences and is highlighted similarly. (C) 3Di structural tree estimated using the Q.3Di.AF model. Sequences assigned to the early transfer from the archaeal clade to bacteria are highlight as in (B), but now this transfer is inferred for both the catalytic and non-catalytic subunits. Numbers in red, black indicate branch lengths and ultrafast bootstrap supports respectively. In both cases grey clades represent eukaryotes. The green circles and orange squares indicate cyanobacterial and proteobacterial contributions in eukaryotes representing the plastid and mitochondrial ATPases.

As before, we predicted AlphaFold structures for all 1520 sequences and extracted the 3Di sequences using FoldSeek and calculated a 3Di (structural) ML tree with the Q.3Di.AF as the substitution model. While the tree in this case looks remarkably like the amino acid ML tree, the 3Di structural ML tree has significantly shorter branches (Figure 3C). This new topology also reconfirms the root of ToL as between the archaea and bacteria. Furthermore, in the 3Di tree the early transfer of the archaeal ATPase subunits is recovered basal in both catalytic and the non-catalytic subtrees suggesting a single early transfer from archaea to bacteria. Together with the Elongation factors, our results bolster support for the two-domain tree of life with the eukaryotes branching within archaea. In both these cases it is evident that structural phylogenetics can resolve deep phylogenies and recover consistent groupings within the paralogs despite large divergences in amino acid sequences.

### Evolution of photosystems RCI and RCII

The issue of saturation is not exclusive to tree-of-life problems but to all evolutionarily divergent proteins that share remote homology in sequence. The origin of oxygenic photosynthesis is another event that impacted the overall geochemistry of the planet and has been the subject of contentious debate. Photosynthesis can be classified into two major types: anoxygenic photosynthesis, which uses either reaction centre II (RCII) or reaction centre I (RCI), but never both together and oxygenic photosynthesis which uses both reaction centres I and II (RCI and RCII) coupled to a water splitting reaction that leads to the formation of oxygen (Hohmann-Marriott and Blankenship, 2011). One set of theories suggests that anoxygenic photosynthesis evolved first and later developed into oxygenic photosynthesis (Martin et al., 2018). An alternative view favours oxygenic photosynthesis to have evolved first, with anoxygenic phototrophs having lost either RCI or RCII. One piece of evidence for the latter view is the lack of any bacterial group that harbours the anoxic versions of both RCI and RCII, which is thought to be a necessary precursor to oxygenic photosynthesis (Sánchez-Baracaldo and Cardona, 2020). Until recently, members of the *chloroflexota* phylum have only been known to harbour anoxic RCII. This changed when a *chloroflexota* group, *Ca. Chloroheliales,* was identified that contains RC1 (Tsuji et al., 2024). This still falls short of proving that anoxic RCI and RCII have existed together in the same genome however, one possible interpretation of these data is that an ancestral *Chloroflexus* might have contained both, leading to differential losses in extant lineages of *Chloroflexi*. This would support the idea that anoxic photosynthesis may have come first, if these photosystems are close relatives of the photosystems that were eventually transferred into cyanobacteria The phylogenetic tree based on amino acids of RCI containing *Chloroflexi* does not place their RCI sequences as close relatives to those of cyanobacteria (4A, re-inferred for this study). But this tree suffers from extremely long branches, and we wondered whether this placement is the result of long branch attraction. We therefore set out to re-infer this tree using 3Di characters and our structural substitution matrix (Figure 4B). This shortened all relevant internal branches to lengths well below one but yielded the same topology as the amino acid tree. This confirms the authors’ original inferences and leaves the evolution of oxygenic photosynthesis an unsolved problem for now.

**Figure 4:**
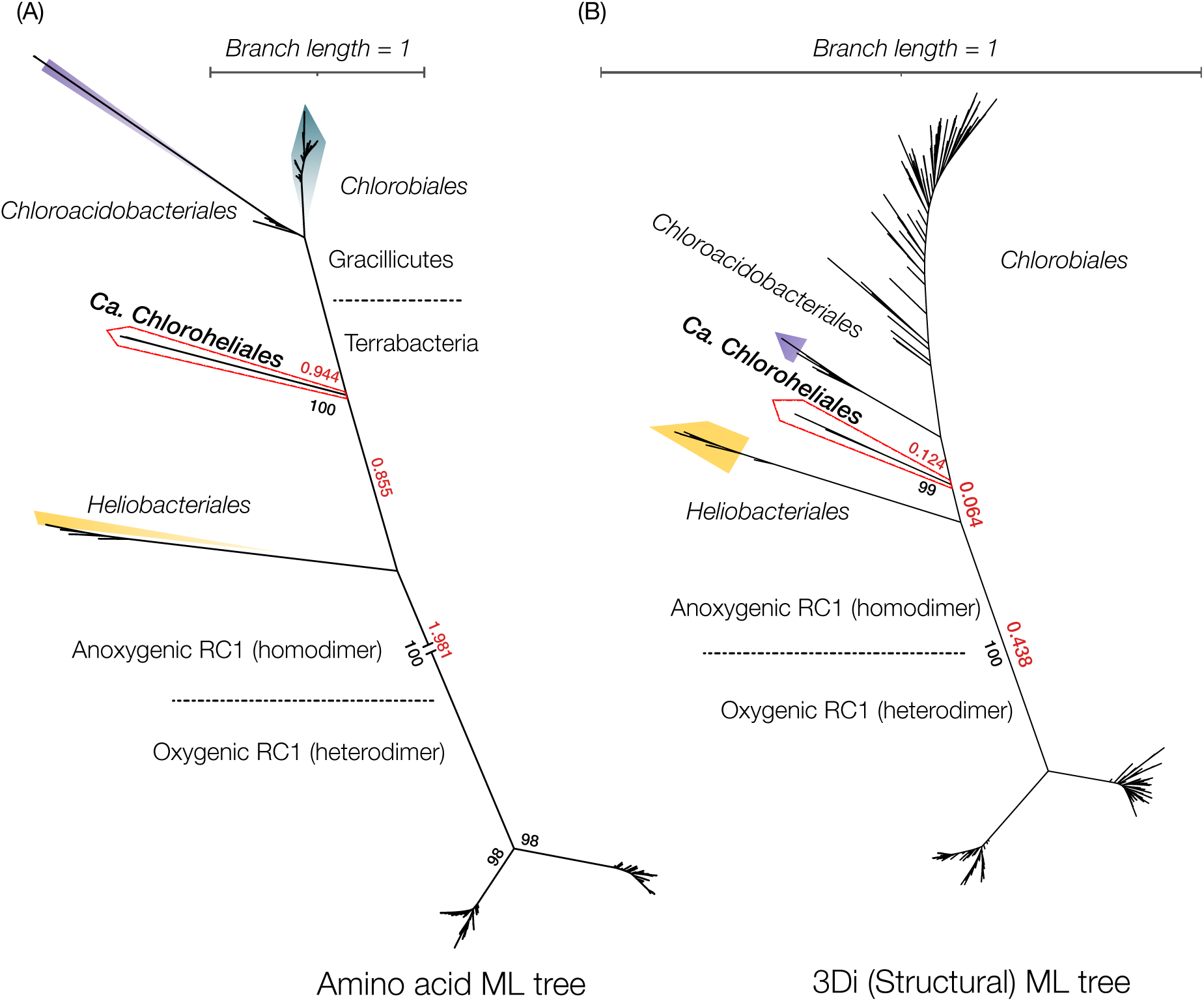
*(A)* Amino acid ML tree of 321 RC1 protein sequences. Note that long branches are broken as indicated for illustration. *(B)* 3Di structural ML tree of 297 3Di sequences from AlphaFold structures using the Q.3Di.AF model. Numbers in red, black indicate branch lengths and ultrafast bootstrap supports respectively.

**Figure 5:**
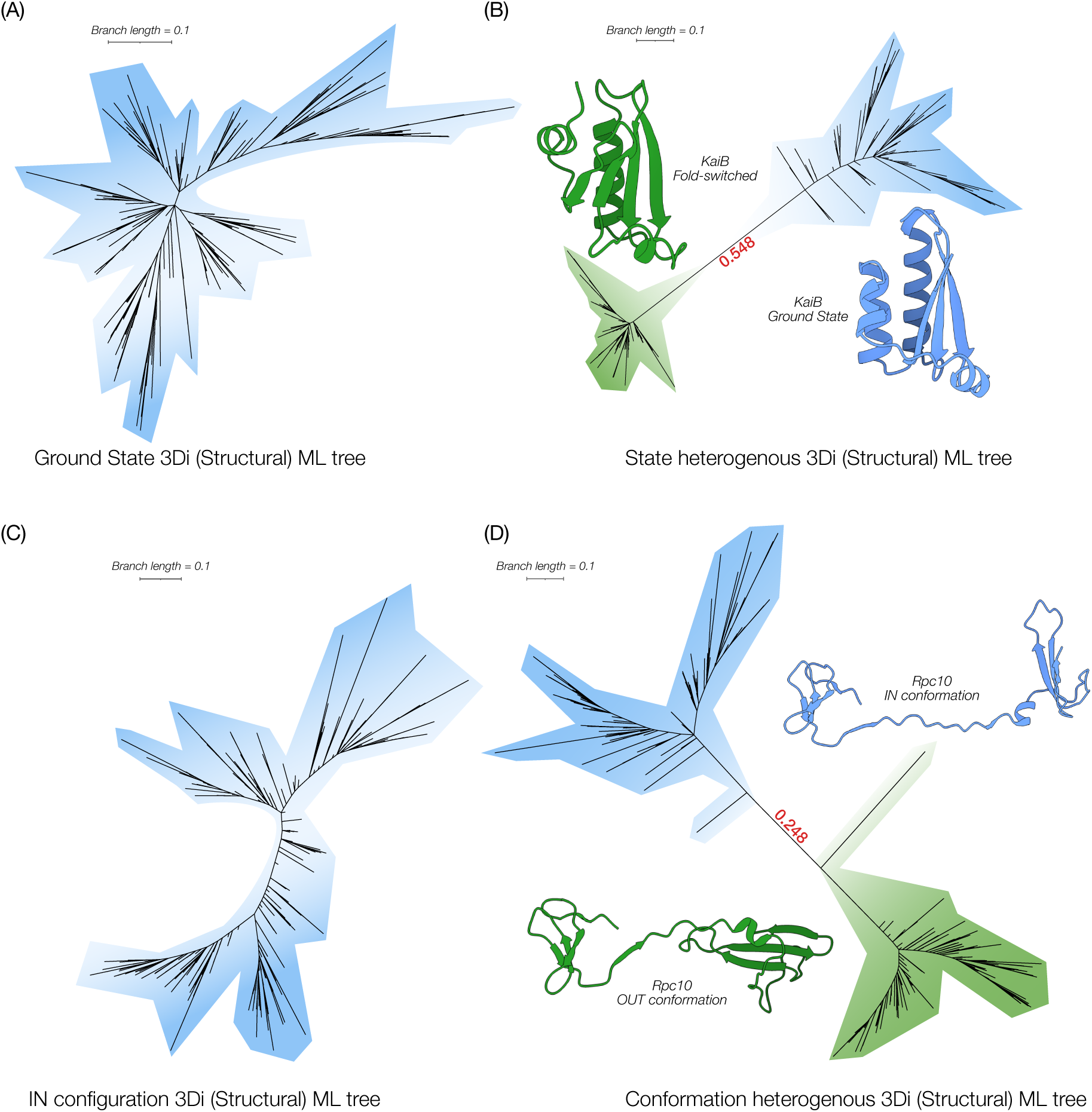
(A) 3Di structural ML tree constructed from KaiB proteins modelled in the ground state. (B) 3Di structural ML tree constructed from approximately 50% of the KaiB proteins modelled in the ground state (blue) and the other 50% modelled in the fold switched state (green). (C) 3Di structural ML tree constructed from RPC10 proteins modelled in the IN conformation (D) 3Di structural ML tree constructed from approximately 50% of the RPC10 proteins modelled in the OUT conformation (blue) and the other 50% modelled in the IN conformation (green). In both cases the distinct conformations form monophyletic groups in contrast to their placements in (A) and (C) respectively.

Our work in this manuscript and that of others (Moi et al., 2023; Puente-Lelievre et al., 2024) clearly points to the utility of structural phylogenetics in cases where structures can be predicted reliably and with one possible structure per sequence. There are several practical and conceptual caveats that come with using this method, which we will briefly elaborate on. We present these caveats in the spirit of critical optimism about the utility and impact of this new method.

### Prediction accuracy of LLMs

Structural phylogenies can only ever be as good as the predicted models that are used to derive 3Di sequences. Predicting large numbers of sequences with AlphaFold is computationally costly and potentially prohibitive for many interested users. Using bilingual Protein LLMs like ProtT5 may seem like an obvious solution, because it removes the computationally expensive requirement of predicting the AF structures of a large number of protein clusters not only in the Q-matrix estimation, but also for tree inference of single protein families with a lot of members. Encouragingly, the Q-matrix estimated from 3Di sequences derived from AlphaFold structures (Q.3di.AF) is very similar to the one from PFAM clusters translated using ProtT5 (Q.3Di.LLM) despite their low accuracy compared to AlphaFold predictions (Figure 1C,1D, Supplementary Figure 1). This could be due to the fact that the LLM has issues when dealing with long repeated stretches which in some cases leads to possible register shifts of structural motifs (Supplementary Figure 2). These register shifts can be dealt with during a 3Di sequence alignment using the 3Di scoring matrix, which we did not perform for our pairwise identity calculations (as the two sequences are the same length). It is also possible that prediction errors average out when inferring a Q-matrix for thousands of protein families, even if there are substantial errors in the alignments of any one family. It is clear, however, that ProtT5 translations are not reliable for inferring individual trees. We tested this by using ProtT5 derived 3Di alignments for the three protein families we investigated here. In two out of three cases we recovered phylogenies that either were biologically improbable (Supplementary Figure 4) and/or erroneous with non-sensical topologies (Supplementary Figure 5). Most of these issues stem from the faulty prediction of 3Di sequences. While we did not observe this problem here when using AlphaFold structures, we expect similar issues when using structures that are not confidently predicted by AlphaFold. For now, reliable tree inference only seems possible using AlphaFold generated structures and therefore comes with a significant computational overhead. Better language and structure prediction models are certain to be available in the future and they should make structural phylogenetics more widely accessible.

### Fold-switching and conformational variability

Many proteins undergo conformational changes and some even switch their folds entirely as part of their functions (Chang et al., 2015). Previous analyses using AlphaFold suggests that it can sometimes predict structures in different conformations despite having a strong bias towards one dominant conformation (Chakravarty and Porter 2022; Sala et al., 2023; Wayment-Steele et al., 2023). Since this can lead to different 3Di sequences for the same protein, depending on which conformation it is predicted in, we wondered if this could lead to spurious grouping according to conformation rather than genealogical relationships on 3Di phylogenies. We examined two proteins for this purpose. One is KaiB, which is known to fold-switch as part of its catalytic cycle, involving a drastic change of a helix to a beta-sheet (Chang et al., 2015; Zhang et al., 2023). The other is the RNA Polymerase III subunit Rpc10, which undergoes a conformational change during its function in gene transcription (Girbig et al., 2021).

To test how much this issue can affect 3Di trees, we constructed a worst-case scenario for both proteins. In both cases, we modelled each sequence on the two distinct conformations using homology modelling and inferred their 3Di sequences using FoldSeek. For tree inference, we then randomly chose the 3Di sequence of one of the two possible conformations for each protein, such that approximately half our sequences were predicted in one conformation, and the other half in the other conformation. For both KaiB and Rpc10 we found that the phylogenetic tree splits the two conformational states with long branches (Figure 4B, 4D) as opposed to a 3Di structural tree which was inferred from 3Di sequences reflecting a single conformation (Figure 4A, 4C). This highlights a severe limitation of structural phylogenetics where the presence of multiple predicted conformations can generate spurious branches and relationships. Here we concocted an extreme case by forcing sequences randomly into distinct conformations. However, in cases where only a small minority of proteins within the analyses share a different conformation, these artefacts can lead to false conclusions. It is therefore very important to assess the conformational homogeneity of the predicted sequences before inferring a 3Di tree.

### Site-independence in structural alignments

One of the main assumptions of maximum likelihood is site independence, which allows the likelihood to be computed independently for all sites in the alignment (Liò and Goldman 1998). It has been obvious for a long time that this is not a realistic assumption. Epistasis between amino acids is a well demonstrated phenomenon and quite extensive among proteins (Starr and Thornton 2016). This violates the site-independence assumption of maximum likelihood phylogenetics, even though it has been argued that increasing the number of sites normally associated with a protein sequence or increasing the number of proteins used for a concatenated alignment averages out the signal in most cases (Starr and Thornton, 2016; Magee et al., 2021). In the case of the structural phylogenetics and 3Di alphabet however, this assumption is explicitly violated since each letter corresponds to at least 6 other amino acid positions in 3D space. It is for example not clear to us that it is even possible for a single substitution to occur at the level of 3Di characters, because of the structural dependence between sites. In a sense, structural phylogenetics makes the ugly truth of model violation explicit in its alphabet. Whether or not this approach becomes widely accepted in evolutionary biology will depend on investigating the consequences of this violation, which is beyond the scope of this manuscript.

### Information loss

The 3Di alphabet compresses information that would be present in amino acids. This is the very reason for its utility in deep phylogenetics, because it overcomes the saturation problem. But it also makes evolution on short time-scales is harder to resolve using these models, and relationships at the very tips of 3Di trees probably much less reliable than in an amino acid or DNA tree (Mutti et al.,2024). A potential solution is to use partitioned models, in which a tree is inferred from both 3Di and amino acid alignments simultaneously, using different substitution models for the partitions (Puente-Lelievre et al., 2024). To make this approach work, however, one would have to allow the structural partition to also have a different set of branch lengths than the amino acid partition (Lopez et al., 2002), which the first use of this approach did not yet include (Puente-Lelievre et al., 2024). Such a heterotachous model presents a difficult optimization problem, which in our hands leads to impractically long run times on our datasets. Another question is the size of the alphabet. 3Di uses 20 characters because this allows simple integration with existing phylogenetic software. It is not yet clear that whether this is even close to the optimal number of characters for structural phylogenetics. Larger alphabets could perhaps retain more short-term information. They would, however, make the inference of substitution matrices much harder.

### Structural phylogenetics and the future of deep history

Our work complements and builds on other recent tools that utilise the 3Di alphabet for structural phylogenetics (Moi et al., 2023; Puente-Lelievre et al., 2024). Our structural Q-matrices should make it much easier to infer structure-based trees for anyone familiar with maximum likelihood phylogenetics. Newly developed online tools for the generation of 3Di alignments should further lower the bar for adoption (Gilchrist et al., 2024). As with every new method, it is difficult to know exactly what impact structural phylogenetics will have. For now, we see its main utility in solving difficult rooting problems involving distant outgroups that amino acid phylogenies cannot solve with any degree of confidence. This will help polarize the direction of evolutionary change in the emergence of many important functions. Better resolved deep protein phylogenies will also improve our reconstructions of the gene content of ancient organisms (The Last Universal Common Ancestor, the Last Eukaryotic Common Ancestor, and the Last Archaeal Common Ancestor, for example).

For now, these methods will not be useful for ancestral sequence reconstruction, because 3Di sequences cannot be back translated into a unique amino acid sequence (Heinzinger et al., 2023). Even though our matrix allows us to infer 3Di sequences at internal nodes of structural phylogenies, it is at present not possible to then turn these reconstructed 3Di sequences into resurrected proteins composed of amino acids. It may, however, be possible to restrict a set of plausible amino acid reconstructions at one particular node on an amino acid phylogeny to a subset that agrees with the reconstructed ancestral 3Di sequence at the corresponding node on a structural phylogeny.

The true impact of viewing the past through the glacial change in the structure of proteins will only emerge when this method is robustly tested and becomes widely adopted in evolutionary biology. We hope the matrix inferred here will be a first step in this process.

## Methods

### Datasets for QMaker

The SwissProt AlphaFold database (https://alphafold.ebi.ac.uk/download) was downloaded and then clustered with FoldSeek (https://github.com/steineggerlab/foldseek) *easy-cluster* with default settings and a coverage of 80%. This yielded 1660 clusters which contained at least 50 members and a maximum of 2500 members. Databases of PDB structures were then created and 3Di sequences were subsequently extracted from these 1660 clusters using FoldSeek as previously described. The PFAM sets were taken from Minh et al., 2021 which contained 6655 protein families used for training the Q-matrix and a further 6653 families were used for testing. In the case of PFAM families the amino acid FASTA files were directly translated to the 3Di alphabet using the scripts provided by Heinzinger et al.,, 2023 (https://github.com/mheinzinger/ProstT5).

### Q-matrix estimation

Both the AF-db and PFAM-db sets of 3Di sequnces were aligned using *ginsi* method within Mafft (v7.515) and the 3Di scoring matrix from FoldSeek using the *–-aamatrix* flag implemented within mafft. The 3Di MSAs thus generated were then used in the QMaker routine as described in Minh et al.,, 2021 (http://www.iqtree.org/doc/Estimating-amino-acid-substitution-models). Briefly, for each MSA the best fit substitution model was initialised with GTR20 along with the best RHAS model to account for rate-heterogeneity using ModelFinder (Kalyaanamoorthy et al., 2017). In the Next step we estimate a joint reversible Q-matrix for all the 3Di MSAs as described.

### Individual Protein/3Di sequences and Phylogenetic tree reconstructions

Elongation factors and Reaction Centre I homologs were identified using BLAST against the NCBI non-redundant (*nr*) database, and then filtered with a minimum similarity threshold of 50% and an e-value cutoff of 1E-5. For the ATPase phylogeny was exactly reproduced from Mahendrarajah et al.,, 2023 and the same sequences used for the 3Di sequences. The amino acid sequences were aligned using *linsi* and then subsequently trimmed using TrimAl (v1.4) (Capella-Gutiérrez et al., 2009) with the *-automated1* setting. Trimmed amino acid alignments were then used for maximum likelihood tree estimation using IQ-tree with the best-fit model suggested by ModelFinder. 3Di sequences for individual proteins trees were extracted from PDB files from individual AlphaFold (v2.2.0) predictions. The best ranked AlphaFold models were used to create a database using FoldSeek which allowed us to extract 3Di sequences from the PDB structures. For 3Di sequences translated from ProtT5, the model was queried as described in Heinzinger et al., 2023 using amino acid sequences as input. All 3Di sequences were aligned with Mafft (*ginsi*) using the --aamatrix option specifying the 3Di scoring matrix provided by van Kempen et al., as part of FoldSeek. 3Di MSAs were then used to estimate the structural ML tree as described above. For individual 3Di tree reconstructions IQ-tree (v2.3.0) was used to identify the best-fit model (Q.3Di.AF, Q.3Di.LLM or GTR20) according to AICc, along with rate-heterogeneity using ModelFinder. Both amino acid and 3Di trees were estimated with 10000 Ultrafast bootstraps (-bb) and 10000 iterations for SH-test (-alrt).

### Homology Modelling

For the KaiB and RPC10 proteins homologs were identified via BLAST as described above. Then PDB structures or AlphaFold structures of the two conformations in question were used as a template in SWISS-MODEL (Waterhouse et al., 2018). KaiB was modelled using the PDB structure 2QKE in the ground state and 5JYT in the fold-switched state from *Thermosynechococcus elongatus*. The RPC10 was homology modelled on the PdB structure 7AE1 in the OUT conformation and 7AE3 in the IN conformation as described in (Girbig et al., 2021). 3Di sequences were extracted from both sets of states/conformations and then randomly sampled to generate a set composed approximately 50% of 3Di sequences from PDB of KaiB and RPC10 in one of the two states/conformation. ML trees were then estimated using these proteins sequences as described above.

**Table 1:** Number of trees that preferred each model/Q-matrix as identified using *modelfinder* from IQ-Tree according to corrected Akaike Information Criterion (AICc), Akaike Information Criterion (AIC) and Bayesian Information Criterion (BIC)

| Model | AICc | AIC | BIC |
| --- | --- | --- | --- |
| Q.3Di.AF | 2342 | 2065 | 2309 |
| Q.3Di.LLM | 3925 | 2958 | 3697 |
| 3DiPhy | 278 | 322 | 267 |
| GTR20 | 108 | 1308 | 380 |
| **Total** | **6653** | **6653** | **6653** |

## Supporting information

Q-Matrix from AlphaFold predictions

Q-Matrix from ProtT5 LLM predictions

## Supplementary Figure Legends

**Supplementary Figure 1:**
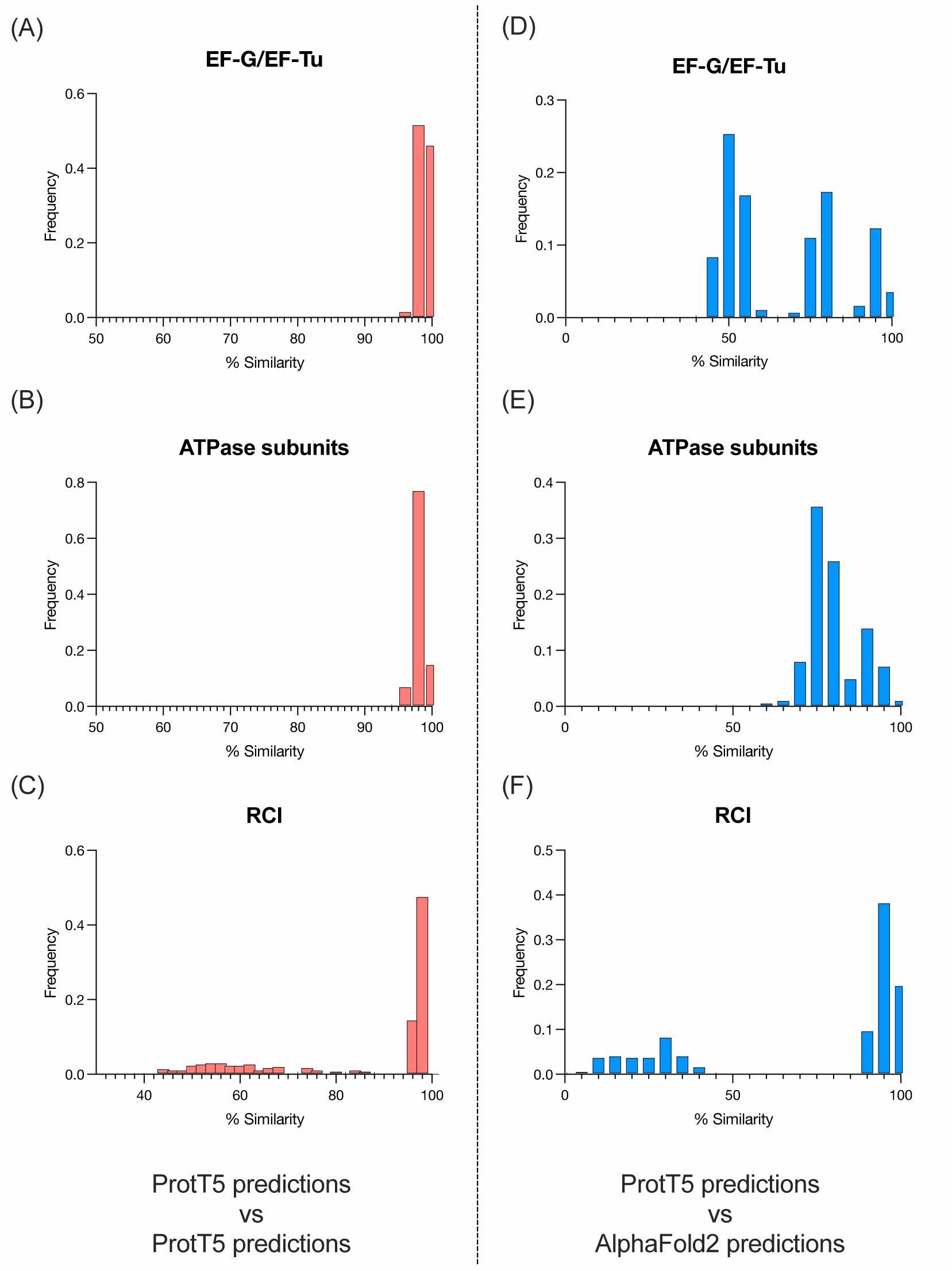
(A-C) Average Percentage similarity between 10 independent rounds of 3Di translations using the ProtT5 model for Elongation factors, ATPase subunits and the Reaction Centre I proteins respectively. (D-F) Percentage similarity between 3Di translation using the ProtT5 model and 3Di sequences extracted from AlphaFold predicted structures for Elongation factors, ATPase subunits and the Reaction Centre I proteins respectively. In all cases percentage similarities were calculated based on the BLOSUM style 3Di scoring matrix on unaligned sequences. Results show that the ProtT5 model is more precise than it is accurate when compared to AlphaFold predictions in all the three cases tested

**Supplementary Figure 2:**
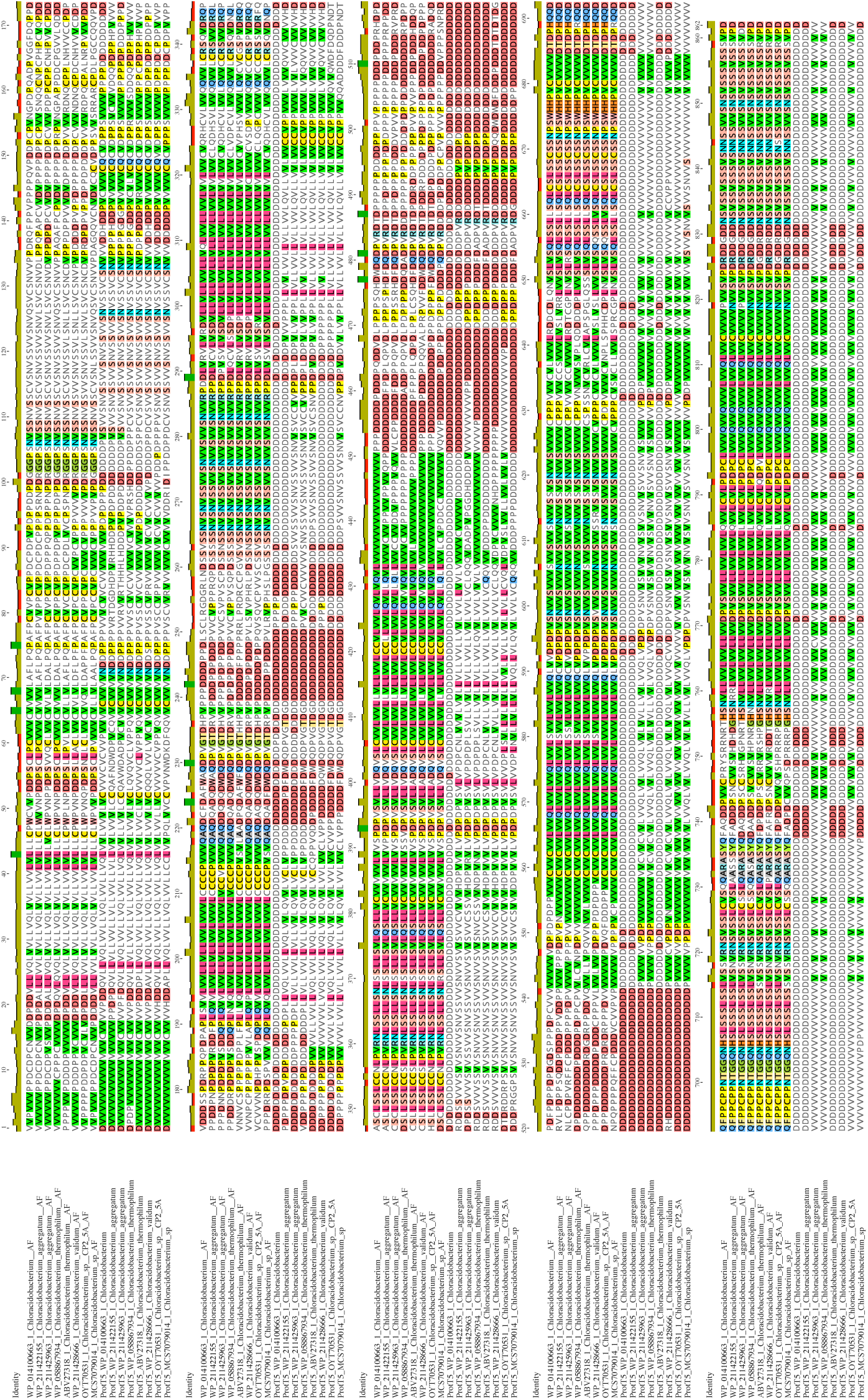
(A) Sequences shown are 3Di translations 549 of representative RC1 proteins prefixed with “ProtT5_” while the 3Di sequences extracted from AlphaFold structures are suffixed with “_AF”. The first observation is the obvious dissimilarities between the ProtT5 (LLM) predictions and the AF extractions. The second observation is that in some cases this erroneous insertions of stretches of 3Di characters are responsible for mismatches observed downstream. This suggests that while the predictions of the ProtT5 LLM is wrong, an alignment guided using the 3Di scoring matrix will be able to align portions of the predictions. This explains the remarkable similarities between the Q557 matrices estimated from the AlphaFold predictions and the LLM.

**Supplementary Figure 3:**
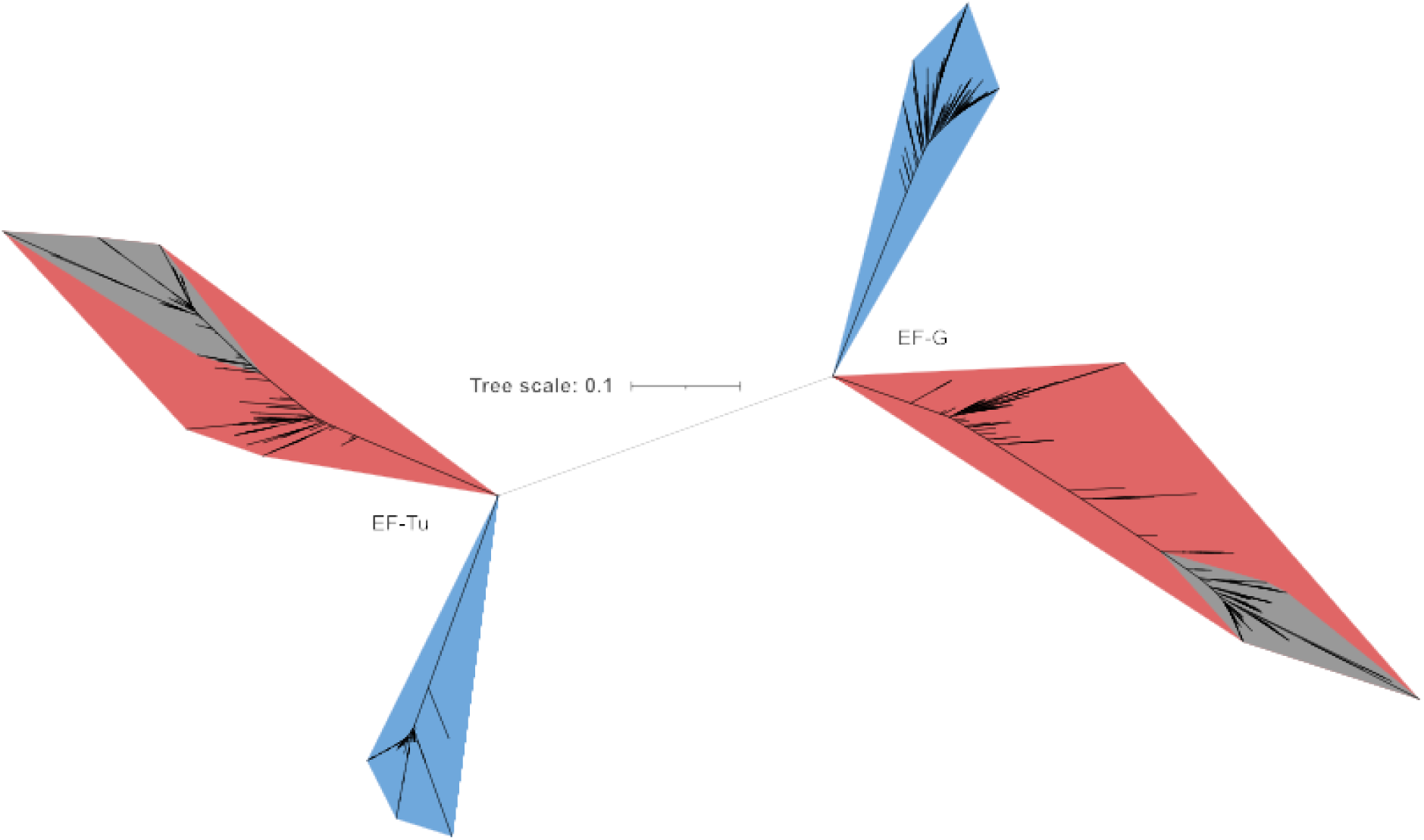
(A) 3Di (structural) ML tree on 3Di translations using ProtT5 of Elongation factor proteins. Red, Blue, and Grey represent Archaeal, Bacterial, and Eukaryotic groups respectively. This particular tree recovers the two-domain topology for the tree of life albeit consistent with the 3Di (structural) ML tree estimated from 3Di sequences extracted from AlphaFold structures.

**Supplementary Figure 4:**
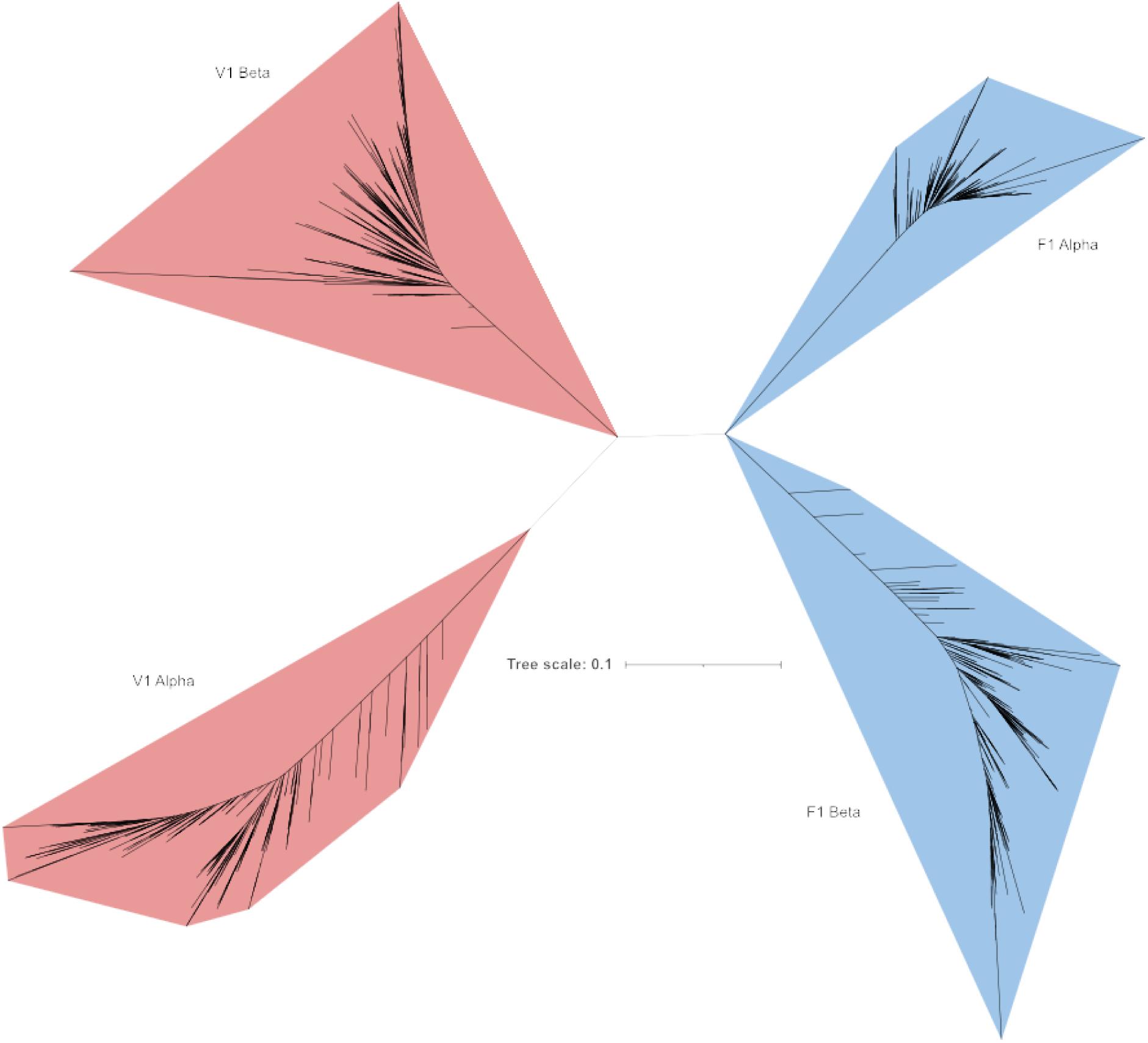
(A) 3Di (structural) ML tree on 3Di translations using ProtT5 of ATPase subunits. Red Blue, and Grey represent Archaeal, Bacterial, and Eukaryotic groups respectively. V1 Alpha and F1 Beta are the catalytic subunits while V1 Beta and F1 Alpha are non-catalytic. This tree recovers a root for the tree of life between archaea and bacteria. It does, however, groups the respective catalytic and non-catalytic subunits of bacteria together, as well as the catalytic and non-catalytic subunits of archaea. This would require an independent loss of catalytic activities in one of the subunits in both the groups. This is inconsistent with currently established theories on the origin of the rotary ATPase. For comparison, our structural tree derived from AlphaFold predictions (Figure 3C) groups archaeal and bacterial catalytic subunits as one monophyletic group and the non-catalytic subunits as another.

**Supplementary Figure 5:**
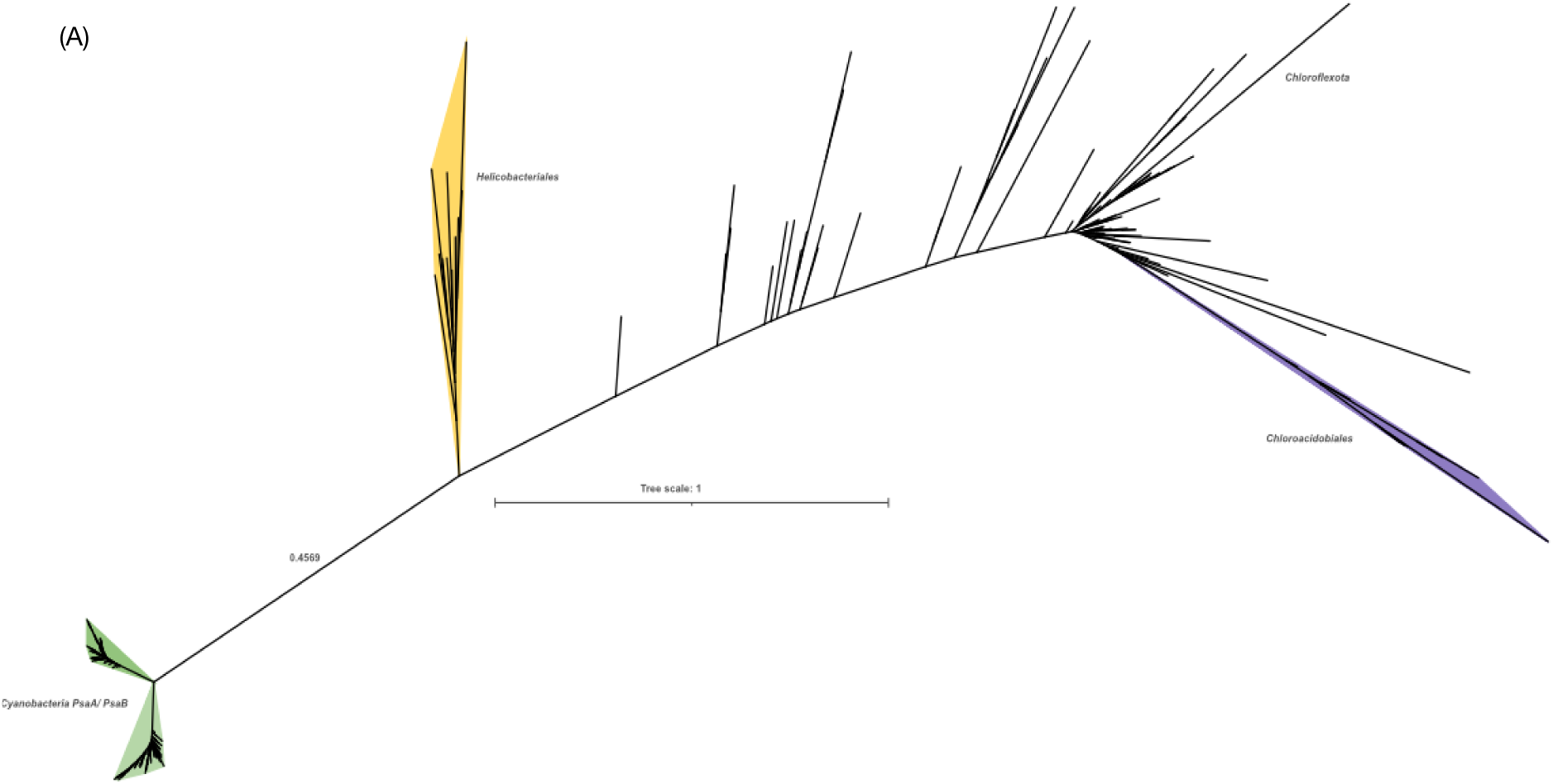
(A) 3Di (structural) ML tree on 3Di translations using ProtT5 of ATPase subunits. This particular tree is highly inconsistent and does not recover the split between chlorobiales and chloroacidobiales. This is also evident from the particularly low similarities between the 3Di translations using ProtT5 and the 3Di sequences extracted form AlphaFold structures. Such cases highlight the importance of the quality of the structure predictions.

## Author Contributions

SGG and GKAH conceptualized, designed and wrote the manuscript. SGG performed the computations

## Acknowledgments and Funding

We would like to thank Mathias Girbrig for suggesting RPC10 for studying the impact of conformational heterogeneity. The study was funded by the Human Frontiers Science Program Grant (RGP0028) awarded to GKAH.

## Data availability

All datasets, trees and alignments are available at https://edmond.mpg.de/privateurl.xhtml?token=624d9e21-f33d-408b-8a81-93d9ad020426 for review and will be made fully public on publication.

